# Exotoxins secreted by *Clostridium septicum* induce macrophage death: *implications for bacterial immune evasion mechanisms at infection sites*

**DOI:** 10.1101/2024.06.18.599648

**Authors:** RM Ortiz Flores, CS Cáceres, TI Cortiñas, SE Gomez Mejiba, CV Sasso, DC Ramirez, MA Mattar Domínguez

**Affiliations:** Department of Human Physiology. School of Medicine. CAMPUS TEATINOS C/Boulevard Luis Pasteur. University of Malaga. 29010-Malaga, Malaga, Spain; Laboratory of Microbiology, School of Chemistry Biochemistry and Pharmacy. National University of San Luis, 5700-San Luis, San Luis, Argentina; Laboratory of Experimental Therapeutics and Nutrition, IMIBIO-SL, CCT-San Luis. CONICET-National University of San Luis. 5700-San Luis, San Luis, Argentina; Department of Medicine and Dermatology, School of Medicine. CAMPUS TEATINOS, C/Boulevard Luis Pasteur. University of Malaga. 29010-Malaga, Malaga, Spain; Laboratory of Experimental and Translational Medicine, IMIBIO-SL, CCT-San Luis. CONICET-National University of San Luis. 5700-San Luis, San Luis, Argentina

**Keywords:** *Clostridium septicum*, exotoxin, partially purified fraction, macrophages, cell death mechanism, apoptosis, autophagy

## Abstract

The induction of macrophage death is regarded as a potential mechanism by which components secreted by *Clostridium septicum* are used to evade the innate immune response and cause tissue damage. This study aimed to determine the effect of partially purified fractions of extracellular proteins secreted by *C. septicum* on the death of mouse peritoneal macrophages. Elicited mouse peritoneal macrophages were incubated with partially purified fractions of proteins secreted by *C. septicum* into the culture medium. After incubation, we found that the protein fraction with a molecular weight ≥ 100 kDa caused significant cell death in macrophages, changed cell morphology, increased markers of apoptosis and autophagy, and increased the expression (protein and mRNA) of IL-10 and TNFα Our data suggest that the proteins secreted by *C. septicum* (MW, ≥ kDa) induce cell death in macrophages by promoting autophagy- triggered apoptosis. This study may contribute to our understanding of the molecular mechanism of immune evasion by *C. septicum* at the infection site.

## 1. Introduction

A surprising feature of myonecrotic disease caused by clostridia is the absence of phagocytic cells at the infection site (Wang and Cull 2022). Circulating monocytes are recruited to the injury site, where they differentiate into tissue macrophages. These macrophages eliminate infectious agents, eliminate damaged cells, and repair tissue (Bryant and Stevens n.d.; O’Brien and Melville 2004). Recent studies suggest that extracellular proteins (as a whole or fractions) secreted by clostridia can cause membrane pores in phagocytic cells and phagosome membranes (O’Brien and Melville 2004; Stevens, Aldape, and Bryant 2012).

Gram-positive bacteria, such as Staphylococcus, Streptococcus, Bacillus, Listeria, and Clostridium, are capable of producing programmed cell death (apoptosis) in eukaryotic cells. One of the secreted virulence factors that can induce apoptosis is the bacterial wall component lipoteichoic acid, although the available data are contradictory (Lotz et al. 2004). Many apoptogenic components in gram-positive bacteria have been identified, including the exotoxin-like-lethal factor *Bacillus anthracis* (Park et al. 2002; Popov et al. 2002), cytosine-like listeriolysin O of *Listeria* monocytogenes (Carrero, Calderon, and Unanue 2004; Guzmán et al. 1996), and hemolysin-like hemolysin β of *Streptococcus agalactiae* (Liu et al. 2004; Ring et al. 2002).

*Clostridium septicum* is an anaerobic gram-positive terminal endospore-forming bacillus (Katlic, Derkac, and Coleman 1981). This bacterium exerts its pathogenic action through different extracellular toxins, leading to death by toxic shock, necrosis, and inflammation at the infection site (Hickey et al. 2008). The cytotoxicity of the alfa toxin of *C. septicum* has been the most investigated in different cellular lines. For example, it has been found that the toxicity mechanism depends on the eukaryotic cell type used to assess cytotoxicity (Kennedy et al. 2009). Indeed, the protoxin form of *C. septicum* α- toxin causes a significant cytolytic effect in eukaryotic cells (Srivastava et al. 2017). Furthermore, this protoxin causes pores in the cell membrane and triggers apoptosis (Kennedy et al. 2009).

Because of the extensive tissue damage and high mortality observed in the nontraumatic gas gangrene 67-100% caused by *C. septicum* both in humans and in animals, it is noteworthy that in addition to the cytolytic activity of α-toxin, other exotoxins secreted by the bacterium may participate in the death of host tissue cells (Dedemadi et al. 2011). However, studies on the identification of *C. septicum* exotoxin/s that cause macrophage death are rare (Srivastava et al. 2017).

Autophagy, also known as programmed cell death type II, is a distinctive cellular event characterized by the formation of autophagosomes and the degradation of organelles and intracellular materials within autophagosomes fused with lysosomes (Klionsky et al. 2012; Yu, Chen, and Tooze 2018). In addition to occurring at a basal level, this process is enhanced in response to stress conditions such as starvation, radiation, or pharmacological agents (Chen et al. 2009). Whether exotoxins secreted by *C. septicum* cause autophagy in macrophages remains a matter of discussion (Mihalache and Simon 2012).

Herein, we aimed to investigate the effects of partially purified fractions (PPFs) of proteins secreted by the culture of *C. septicum* on the mechanism of macrophage death, which may help explain bacterial evasion of the innate immune response at the infection site.

## 2. Material and Methods

### 2.1. Clostridium septicum

The ATCC 12464 strain of *C. septicum* was obtained from the American Type Culture Collection (Manassas, VA, USA). This strain was preserved under anaerobic conditions in a cooked meat culture (MCC) medium at room temperature. An anaerobic environment was achieved using the Vaseline- Paraffin (VAS-PAR) method. Both cell and cell supernatants were obtained when the bacterial culture reached the logarithmic growth phase. The evolution of the cultures was followed by changes in optical density at 580 nm (DO580) using a Metrolab VD 40 spectrophotometer. The early logarithmic phase of the culture was obtained between 4 and 6 h of incubation.

### 2.2. Preparation *C. septicum* partially purified fractions (PPF)

To obtain the cell-free supernatant (CFS), the cultures in the logarithmic growth phase were centrifuged at 5,500 × g for 10 min at 4°C (SIGMA, 3K30 centrifuge). The pelleted cells were then separated from the supernatant. The PPFs were obtained from the CFS by ultracentrifugation fractionation according to the molecular weight cutoff using Millipore 100 kDa and 30 kDa cutoff concentrators and an Amicon Ultra0.5 concentrator of cutoff = 10 kDa at 5,000 × g at 4°C and a recovery spin of 1,000 × g. CFS was concentrated approximately 20 times. PPF was used immediately after being obtained in subsequent tests to avoid any activity loss due to freezing and thawing. PPF was recovered in sterile Eppendorf tubes. The molecular weight (MW) range of the four PPFs used in this study was as follows: (i) ≥ 0 kDa (PPF≥ kDa); (ii) 100 kDa at 30 kDa (PPF100-30 kDa), (iii) ≥ kDa (PPF≥ 0 kDa), and (iv) <10 kDa (PPF<10 kDa).

Isolation and culture of elicited mouse peritoneal macrophages. Elicited peritoneal macrophages were obtained from Rockland mice having between 18 g–20 g of body weight. All protocols for the use of animals were approved by the Institutional Animal Ethical Committee’s (IAEC) guidelines (CICUAL-UNSL-CONICET- CUDAP: BIOL: 2132/2021). Groups of two animals were intraperitoneally injected with 1 ml of a sterile 10% protease peptone suspension (Difco) to increase the number of macrophages in the peritoneum. After 48 h, peritoneal cells were isolated by lavage of the peritoneal cavity twice with 10 ml of ice-cold sterile saline solution (pH 7.4). The cell suspension was centrifuged at 4°C at 3,000 xg for 15 min. The pelleted peritoneal cells were rinsed twice with an ice-cold sterile saline solution. The viability of isolated peritoneal cells was assessed using the trypan blue uptake assay. Isolation of macrophages from the peritoneal cell suspension was performed by selective adherence to a tissue plate in Dulbecco’s modified Eagle’s medium (DMEM) supplemented with 10% fetal bovine serum (FBS), 1% antibiotic-antimycotic mix (Gibco^®^), and 25 μg/ml amphotericin B. After the selection of peritoneal macrophages, the monolayer was rinsed with sterile saline to eliminate nonadherent cells, and fresh medium was added to the macrophage monolayer.

### 2.3. Treatment of macrophages with different PPFs

Macrophages (2×10^6^) were plated in 12- or 24-well plates and incubated for 4 h at 37°C and 5% CO2. After rinsing the monolayers with fresh medium, the macrophages were treated with 1% to 2.5% (%v/v) PPFs in the culture medium for different periods in a cell incubator.

### 2.4. CHO cell culture and autophagy analysis

The hamster-ovarian epithelial cell line CHO-K1 (catalog #CCL-61), stably transfected to overexpress pEGFP-LC3B, was grown in α EM supplemented with 10% FBS, streptomycin (50 μ ml), and penicillin (50 IU/ml). For autophagy analysis, cells were rinsed three times with ice-cold sterile PBS and incubated in Earle’s balanced salt solution at 37°C for different periods in the presence or absence of different concentrations of PPFs. Starvation, a positive control for autophagy, was induced by rapamycin treatment (50 µg/ml). GFP expression in CHO cells was assessed by confocal imaging using an Olympus Confocal FV1000 microscope. Autophagy was assessed by counting, in the confocal images, the number of LC3B dots per cell and by normalizing the results according to the respective control (cells without any treatment). Confocal images from 10 random fields were quantified, representing approximately 80 cells per experiment.

### 2.5. Observation of morphological changes

Morphological changes in macrophages exposed to PPFs were evaluated without staining (40X) or after Giemsa staining. Giemsa staining was performed on 3 mm circular coverslips placed at the bottom of 6-well plates. The cells were treated with PPFs, fixed with methanol, and stained with Giemsa (1/24) for 30 min. The program FV10-ASW 3.0 was used for living-cell image analysis. The observed morphological changes were expressed as the number of apoptotic or necrotic cells observed in 100 cells analyzed.

### 2.6. DNA leader-fragmentation analysis

After the indicated incubation periods with PPFs, macrophage monolayers were rinsed with ice-cold PBS and then lysed with 350 µL of lysis buffer (NaCl (1 M), sucrose, (1 M), Tris (1 M), EDTA (1 M), and SDS (10%), pH 8.6). Cell lysates were centrifuged at 12,000 × g for 10 min. The DNA contained in the supernatant was precipitated by the addition of three volumes of isopropanol and 5 M NaCl, and the mixture was incubated overnight at -20°C. Afterward, the mixture was centrifuged at 12,000 xg for 10 min, and the pellets were resuspended in 750 μ of Tris-EDTA buffer (0.25 M sodium chloride, 0.05 M Tris-HCl (pH 7.5), 20 mM EDTA, sodium dodecyl sulfate (1% SDS) (w/v) and polyvinylpyrrolidone (4% PVP) (w/v), pH 7,6. Then, 10 μ of DNA was mixed with 2.5 μl of sample buffer (0.1% bromophenol blue, 15% glycerol) and resolved by 1% agarose gel electrophoresis (90 V/cm for 1 h). Finally, the gel was stained with 1% GelRed™ (Genbiotech), and images were acquired using a UV-transilluminator.

### 2.7. RNA extraction and reverse transcriptase-polymerase chain reaction (RT]PCR)

RNA isolation and reverse transcriptase-PCR (RTLPCR) were performed to measure the expression of genes encoding inflammatory cytokines upon treatment of mouse peritoneal macrophages with different PPFs. After exposure to PPFs, the culture medium was removed, and the macrophage monolayer was rinsed twice with sterile PBS (pH 7.4). Total RNA was isolated and purified using TRIzol Reagent (Invitrogen, Cat#) as described by the manufacturer. The purified RNA pellet was resuspended in 50 μl of RNAse-free water. mRNA was retrotranscribed to cDNA according to the M-MLV Transcripta protocol (PB-L Productos Bio-Logicos). Amplification of the cDNA fragments was carried out using 20 μl of the reaction mixture (10X reaction buffer, 2.5 μl, 50 mM MgCl2, 0.75 μl, 10 mM dNTPs (dATP, dTTP, dCTP, dGTP), 1 μl; 10 mM forward (F) and reverse (R) oligonucleotides, 1.25 μl of each, 13 μl nuclease-free H2O (added first) and finally 0.25 µl of Taq DNA Pol (5 IU/μl)). Five microliters of cDNA were used for annealing.

The amplification protocols used for each sequence were those recommended by the GenBank NCBI (http://www.ncbi.nlm.nih.gov). The primers used to amplify the resulting cDNAs were as follows: TNFα, F: CCGAGGCAGTCAGATCATCTT and R: AGCTGCCCCTCAGCTTGA; IL-10, F: TTACCTGGAGGAGGTGATGC and R:

GGCTTTGTAGACCCCCTTCT; GAPDH (housekeeping gene), F: ATCACTGCCACCCAGAAGAC and R: GCACGTCAGATCCACAACAG. Samples (10 μl) were mixed with 2.5 μl of sample buffer and separated by agarose gel (1.9%) electrophoresis at 90 V/cm for 1 h. Afterward, the gel was stained with a 1% solution of GelRed™ (Genbiotech), and the bands were visualized and acquired under UV transillumination. The size of the resulting amplicons was determined by comparison with a molecular weight marker (100 bp PB-L^®^ Ladder).

### 2.8. Western blot

Briefly, macrophages were treated with PPFs, lysed with 150 μ native sample buffer per well, and scraped. The cell lysate was boiled for 4-5 min, and 400 μ of the reducing agent DTT (1%, v/v) was added. Samples were separated by 10% SDSL AGE, blotted onto a nitrocellulose membrane, and then immunostained with primary antibodies (anti-Bcl-1, anti-Bax). Immunocomplexes were detected using anti-rabbit IgG-horseradish peroxidase-conjugated antibody. The immunocomplexes were then developed using the LumiPico^®^ ECL Kit following the manufacturer’s instructions (ShineGene, catalog #ZK00902).

### 2.9. Statistics

Data are expressed as the mean values ± standard errors of the means (SE) from three independent experiments. Statistical analyses were performed with GraphPad Prism software using one-way ANOVA and/or Student’s two-tailed t-test followed by Tukey’s comparisons test. A p<0.05 was considered statistically significative.

## 3. Results

### 3.1. C. septicum PPFs kill mouse peritoneal macrophages

To test the effect of PPFs on macrophage viability, a trypan blue assay was performed (**Fig. 1**). After 2 or 4 h of incubation with 2.5% (v/v) PPF<10 kDa in the culture medium, macrophage viability was not affected. However, incubation with 2.5% PPF>10 kDa, 100-30 kDa and < 100 kDa caused approximately 85% cell death after 2 h of incubation and almost 100% cell death after 4 h.

**Fig 1.**
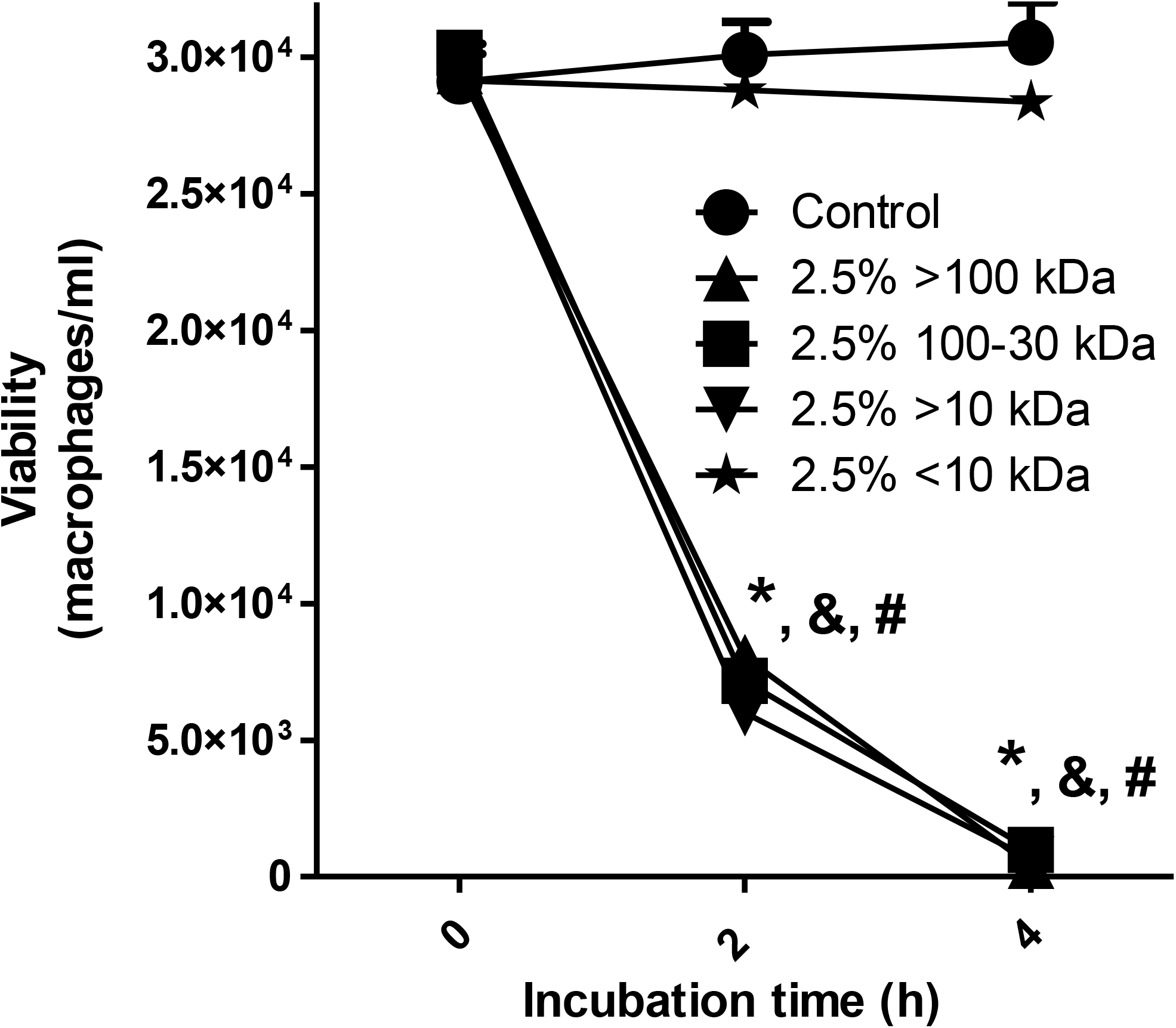
Effect of PPFs on the viability of mouse peritoneal macrophages. Macrophages (3x104) were treated with 2.5% PPFs (≥100 kDa, 100-30 kDa, > 10 kDa, and <10 kDa) in complete culture medium for 4 h. **A**) Cell viability was assessed by the Trypan blue exclusion assay. The controls were cells incubated in complete medium. **B**) Forty to sixty cells were observed in randomly chosen 40X-optical fields per experiment. The results are shown as the mean values ± SEMs from three independent experiments. The signs *, & and # indicate p<0.01 for PPF≥ kDa, 100-30kDa and >10kDa, respectively, compared to the control.

### 3.2. C. septicum PPFs produced morphologic changes compatible with cell death in mouse peritoneal macrophages

To test the mechanism of PPF-induced cytotoxicity caused by PPFs, mouse peritoneal macrophages were treated with 1% or 2.5% PPF ≥ 100-30, ≥ and <10 kDa for 4 h. Morphological changes in macrophages were observed using optical microscopy after Giemsa staining (**Fig. 2A**). A marked apoptotic effect (pycnotic nucleus) was observed upon exposure to 2.5% PPF≥ 00 kDa (**Fig. 2A**, **panels a-b**). We also observed morphological changes compatible with apoptosis upon treatment with PPF100-30 kDa (**Fig. 2A**, **panel c**) at low concentrations and necrosis at high concentrations (**Fig. 2A**, **panel d**). A similar effect was observed upon the treatment of macrophages with PPF<10 kDa (**Fig. 2A**, **panel f**). Treatment with PPF<10 kDa did not show any morphological effects on the cells (**Fig. 2A**, **panels g-h**). Differential cell morphology analysis, as the percentages of normal, apoptotic, and necrotic cells, is shown in **Fig. 2B**.

**Fig 2.**
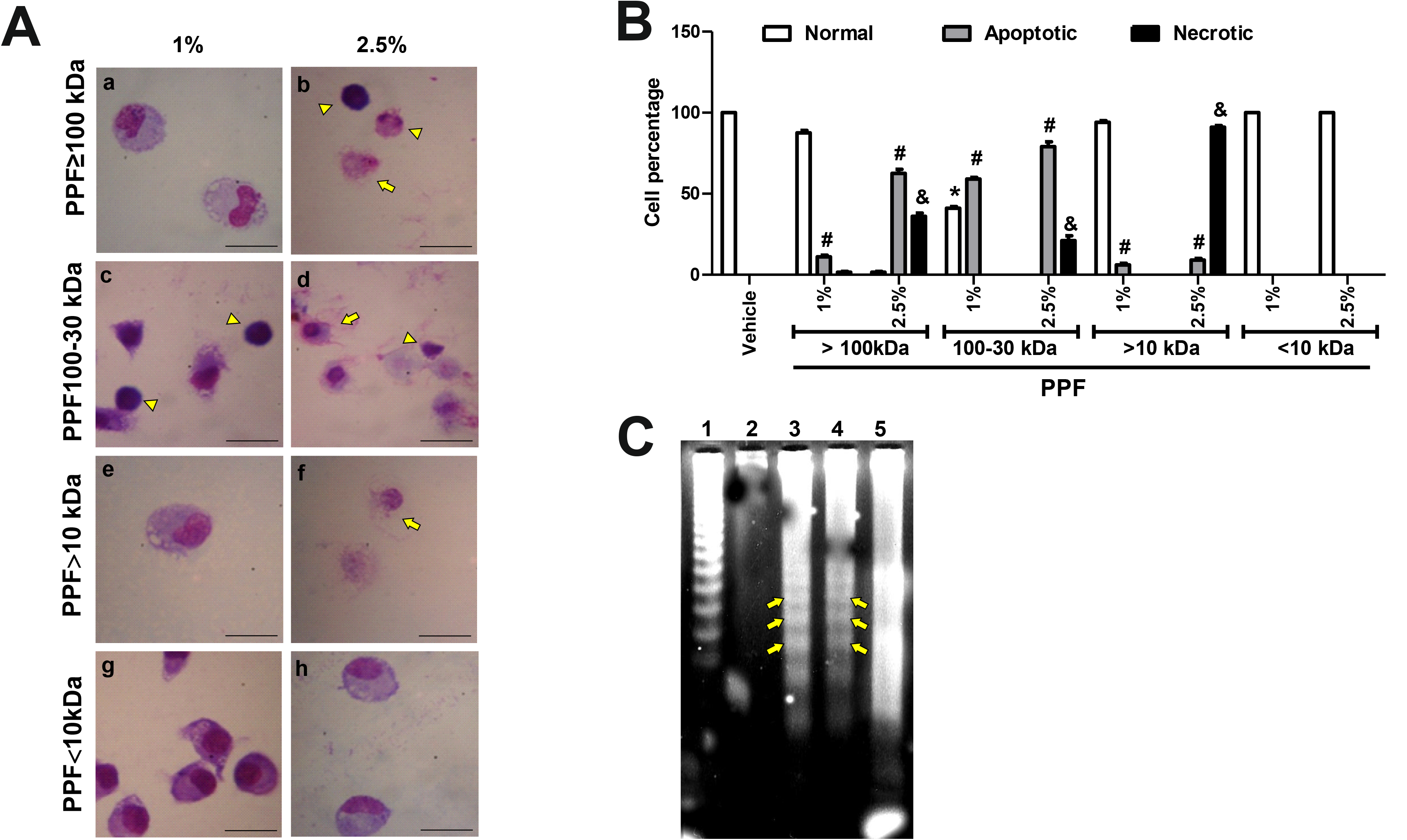
Cell death caused by PPFs. Mouse peritoneal macrophages were incubated with 1 or 2.5% of different PPFs in complete culture medium for 4 h. **A**) 40X micrographs of macrophages stained with Giemsa. Yellow complete arrows indicate morphological changes compatible with apoptosis. Yellow arrowheads indicate cell morphology compatible with necrosis. Measurement bars are 20 μm. **B**) Quantification of cell-morphology changes observed in the micrographs shown in panel A. *, # and & indicate differences (p<0.05) when compared with vehicle alone. **C**) DNA fragmentation analysis in an agarose gel stained by GelRed. The yellow arrows indicate the typical "ladder" pattern corresponding to cell death by apoptosis. Data shown are representative images or mean percent values ± SEM from 3 independent experiments.

An electrophoretic DNA leader pattern is usually observed in cells that die via apoptosis. We observed that treatment of macrophages with 2.5% PPF≥100 kDa resulted in DNA-leader fragmentation (**Fig. 2C**, **Line 3, yellow arrows**). This pattern was not observed in DNA isolated from macrophages treated for 4 h with 1% PPF≥100 kDa (**Fig. 2C**, **line 2**). In contrast, macrophages treated with 1% PPF100-30 kDa showed the typical apoptotic pattern of DNA fragmentation, indicated with yellow arrows in **Fig. 2C**, **Line 4**. However, higher PPF100-30 kDa concentrations in the culture medium caused a DNA-random fragmentation pattern compatible with a necrosis cell death mechanism (**Fig. 2C**, **Line 5**).

### 3.3. C. septicum PPF≥100 kDa induces apoptosis in mouse peritoneal macrophages

To further corroborate the cell death mechanism triggered by PPF≥100 kDa exposure, Bcl-2 and Bax were measured by western blotting using β in as a loading control. Compared to the control, exposure of mouse peritoneal macrophages to 2.5% PPF≥ kDa for 4 h caused a slight increase in the expression of the antiapoptotic protein Bcl-2. However, the same treatment caused an almost 10-fold increase in the expression of the pro-apoptotic protein Bax (**Fig. 3A**). In agreement with these data, semiquantitative analysis of the band intensities shown in **Fig. 3B** indicates an almost 10-fold increase in Bax2 expression.

**Fig 3.**
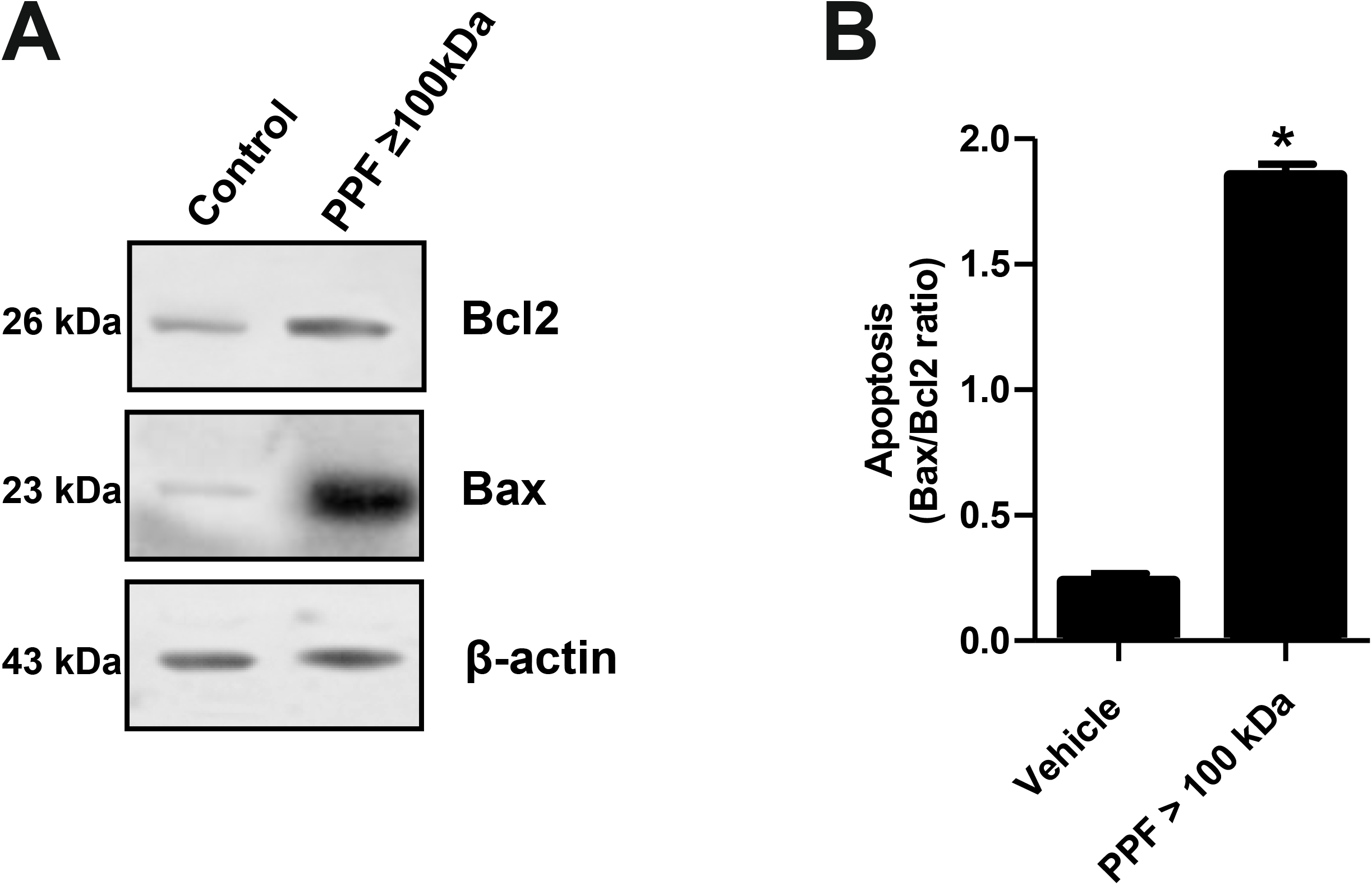
Effect of macrophage exposure to 2.5% PPF ≥ 100 kDa on apoptosis markers. **A**) Western blot of anti-Bcl2, -Bax, and -β -actin (load control). **B**) Bax/Bcl2 band-intensity ratio. Data show a representative image or mean values of density ratios ± SEM from three independent experiments. *p <0.001 compared to the control.

### 3.4. C. septicum PPF≥100 kDa induces autophagy in mouse peritoneal macrophages

Morphologically, macrophages incubated with 1% PPF≥100 kDa showed no significant signs of apoptosis or necrosis but showed extensive development of cytoplasmic vacuoles (see **Fig. 2A**, **panel a**). This morphological pattern raises whether PPF can cause autophagy. Light chain 1 of microtubule-associated protein 1 (LC3B) is the mammalian homolog of the autophagy protein in yeast Atg8, which is added to autophagosome membranes when autophagy occurs. Therefore, we measured the expression of the autophagic protein LC3B in CHO cells stably overexpressing LC-3- green-fluorescent protein (GFP). These cells were incubated for 4 h or 24 h with 0.5%, 1%, or 2.5% PPF≥100 kDa. **Figure 4A** shows a confocal image of the cell localization of LC3B-GFP in cells exposed to different concentrations of PPF≥100 kDa for 4 h and 24 h. An image analysis of **Figure 4A** is shown in **Figure 4B**. The data indicated that the exposure of cells to PPF≥100 kDa caused a higher expression of LC3B-GFP at 4 h than at 24 h of incubation (**Fig. 4B**). Nevertheless, autophagy-induced apoptosis may indicate a possible mechanism for apoptotic cell death triggered by this PPF fraction in mouse peritoneal macrophages.

**Fig 4.**
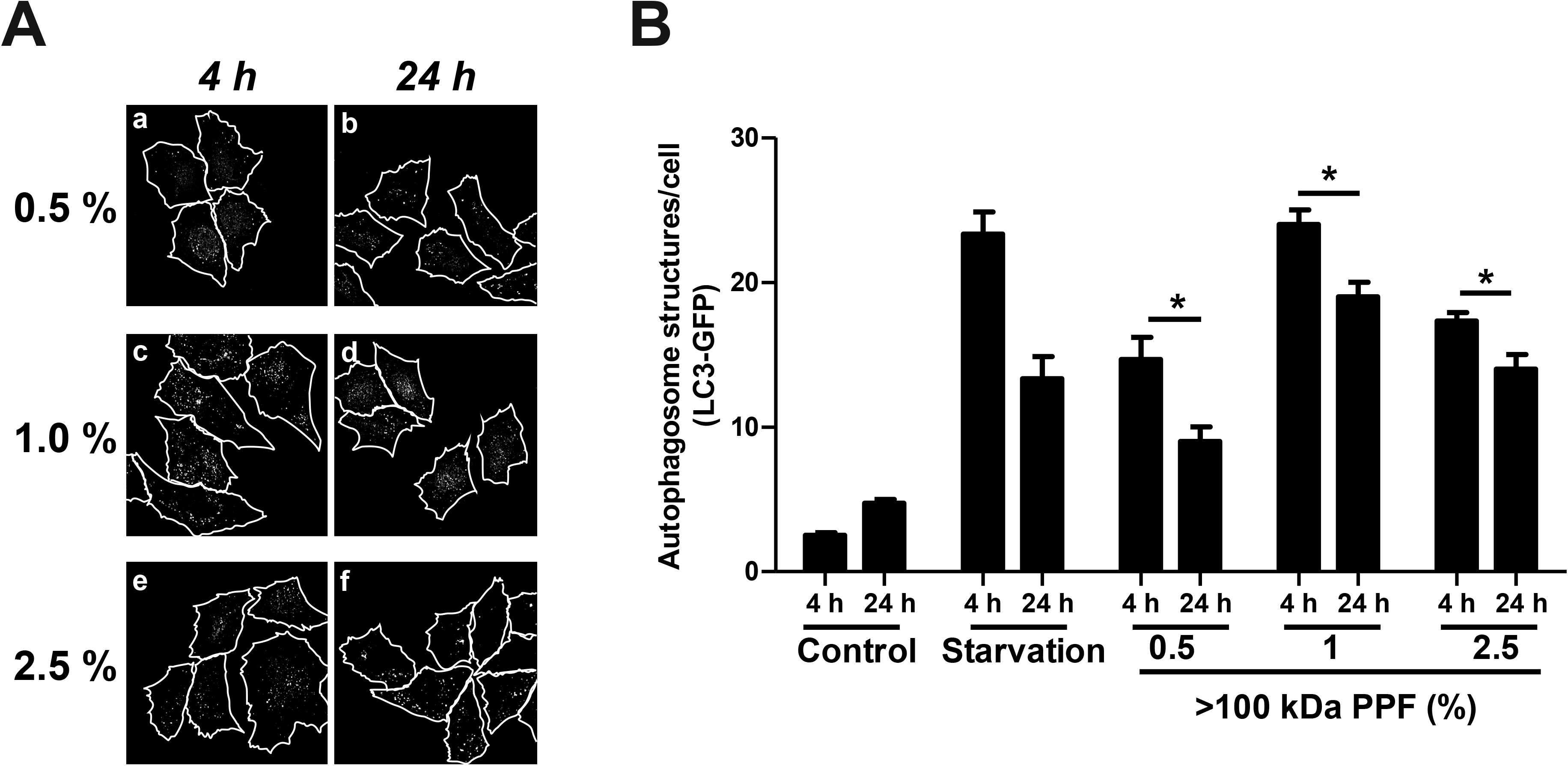
PPF ≥ 100 kDa induces the expression of LC3B, a molecular marker of autophagy. Fluorescence micrograph for the identification of LC3B. **A**) CHO-K1 cells overexpressing pEGFP-LC3B were treated with 0.5%, 1%, or 2.5% PPF ≥100 kDa in complete culture medium for 4 or 24 h. The stitching shows autophagosome structures. B) Quantification of the images shown in panel A. Control-untreated cells; Starvation- cells treated with rapamycin were used as a positive control for autophagy; PPF ≥100 kDa in three different concentrations, i.e., 0.5%, 1.0%, and 2.5%. Data show representative images or mean values ± SEMs from three independent experiments. * p <0.05 when comparing incubation for 4 and 24 h with PPF ≥100 kDa.

To confirm whether autophagy is involved in cell death caused by exposure of mouse peritoneal macrophages to 1% PPF≥100 kDa for 4 h, we measured the conversion of free cytosolic LC3B (LC3B-I) to LC3B conjugated to phosphatidylethanolamine (LC3B-II). This conversion was time dependent, starting from the first hour of exposure to PPF≥100kDa (**Fig. 5A**). The LC3B II/LC3B I ratio, which indicates the conversion rate of LC3B I to LC3B II, increased over time in macrophages incubated with PPF≥ Da 1% (**Fig. 5B**). These data are consistent with autophagy-induced apoptosis and may indicate a possible mechanism for apoptotic cell death triggered by this PPF fraction in mouse peritoneal macrophages.

**Fig 5.**
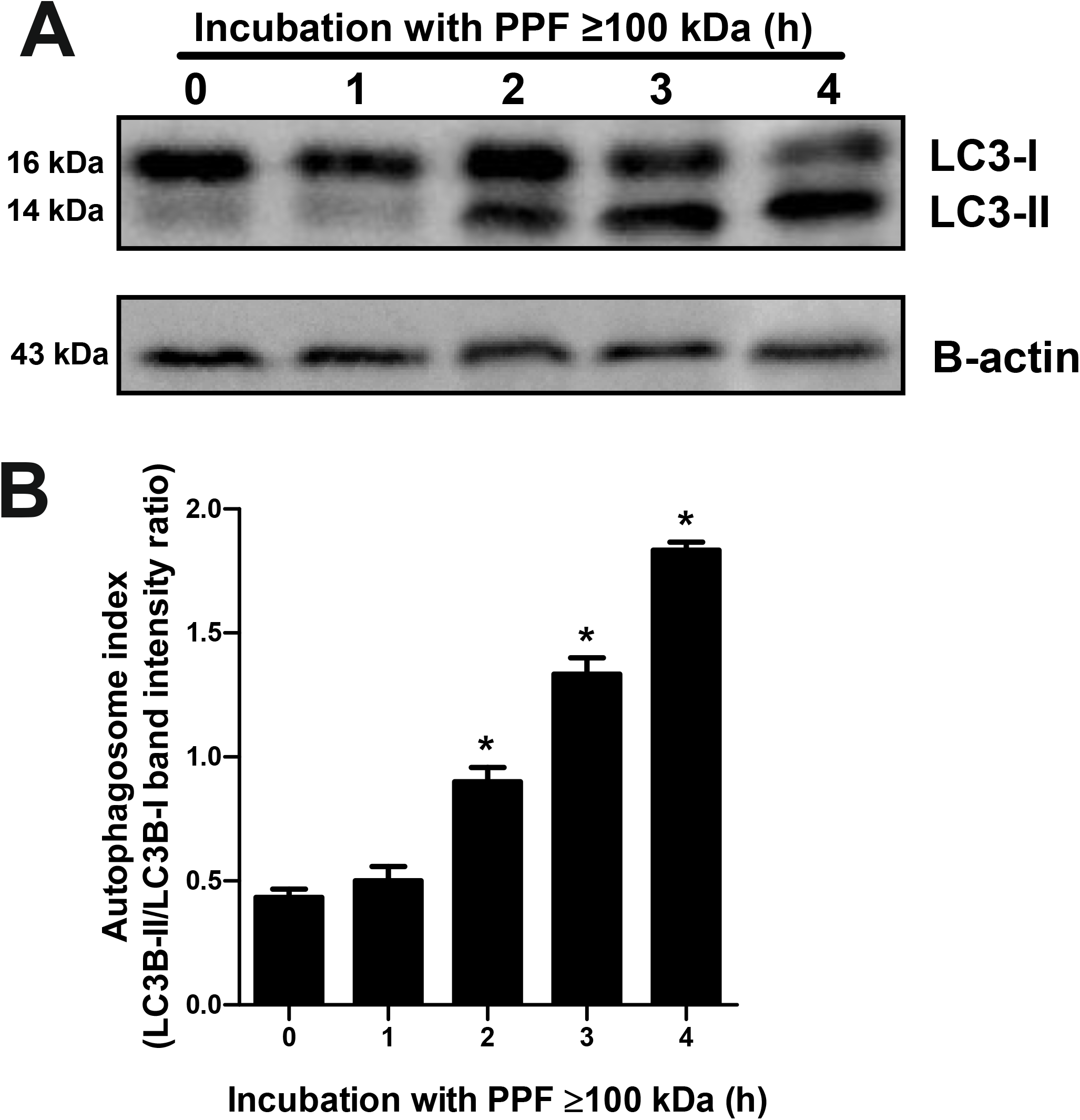
PPF ≥ 100 kDa induces LC3B expression in mouse peritoneal macrophages. **A)** Western blot images showing the conversion of cytosolic LC3B (16 kDa, LC3B-I) into the LC3B-phosphatidylethanolamine conjugate (14 kDa, LC3B-II) after incubation for up to 4 h with 2.5% PPF ≥100 kDa in complete culture medium. β-actin was used as a loading control. **B**) Analysis of band intensities, shown as LC3B-II/LC3B-I intensity ratios at each incubation time. Data are shown as a representative image or as the mean ratio values ± SEM from three independent experiments. * p<0.001 compared to time 0.

### 3.5. C. septicum PPF ≥100 kDa affects cytokine expression in mouse peritoneal macrophages

Several members of the *Clostridium* family secrete proteins that induce the expression of tumor necrosis factor (TNF ) and interleukin-10 (IL-10) in monocytic cell lines. As a cell response to PPF≥100 kDa, the expression of these cytokines may increase. To test our hypothesis, we performed RTL CR analysis of IL-10 and TNFα mRNAs in macrophages exposed to 1 and 2.5% PPF≥100 kDa (**Fig. 6A**). The semiquantitative analysis of band intensities compared to GAPDH mRNA is shown in **Figures 6B**. This result indicates that *C. septicum* PPF≥100kDa can induce the cytokine-mediated response of macrophages, which may be responsible for triggering autophagy and further cell death by apoptosis.

**Fig 6.**
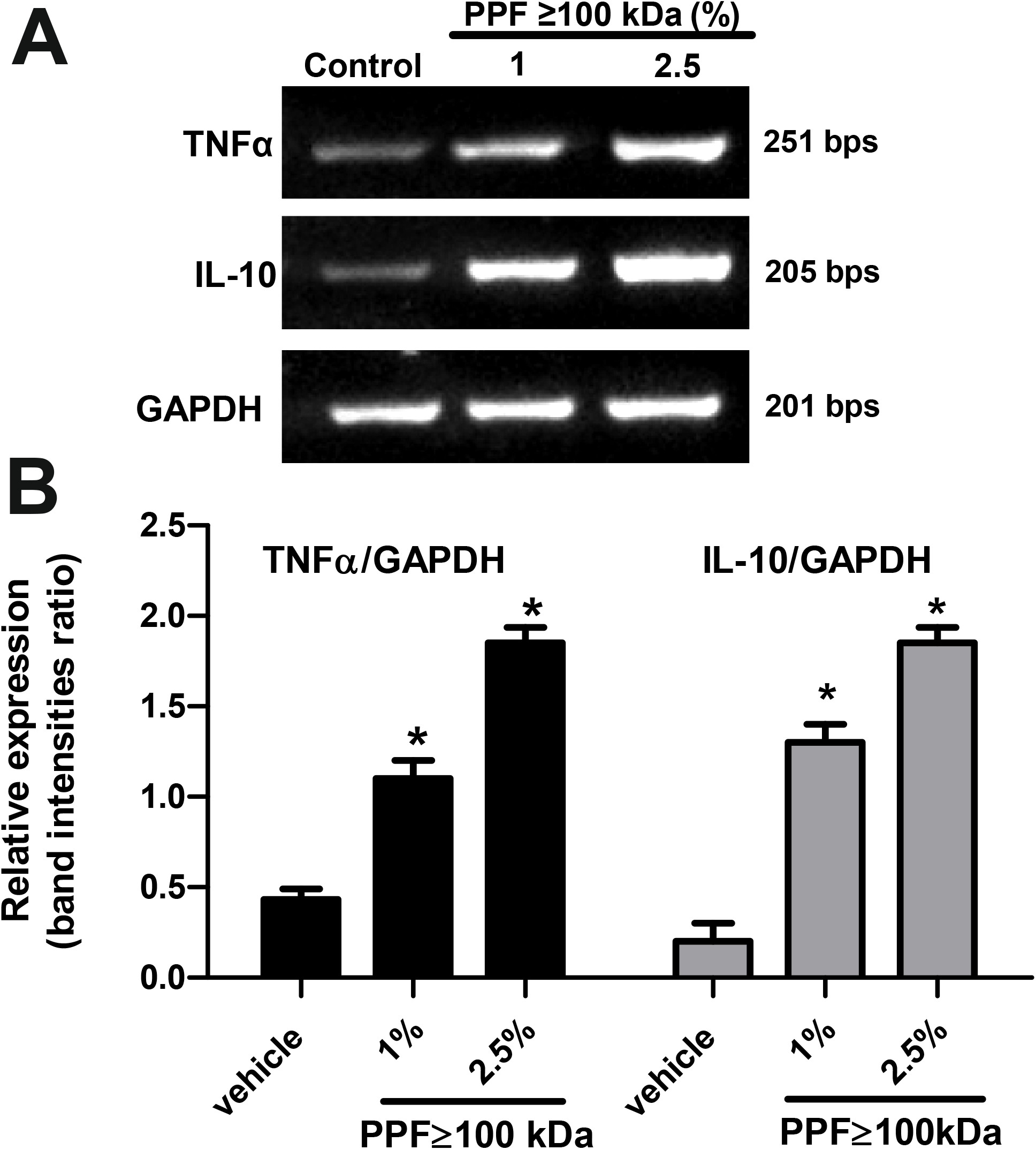
A PPF ≥ 100 kDa induces the expression of TNF-α and IL-10 in mouse peritoneal macrophages.Macrophages were incubated without (control) or with 1% or 2.5% PPF≥100 kDa for 4 h. Then, total RNA was isolated and retrotranscribed, and the resulting cDNA was amplified. **A**)Image showing the resulting RTLPCR amplicons for TNF-α, IL-10, and GAPDH in an agarose gel stained with GelRed. GAPDH was used as a housekeeping gene. **B**) Relative expression of TNF-α and IL-10 shown as TNF-α/GAPDH or IL-10/GAPDH band-intensity ratios. Data show a representative image or mean values of the ratios ± SEM from three independent experiments. *p <0.001 compared to the control.

## 4. Discussion

Exposure of mouse peritoneal macrophages to *C. septicum* PPF≥ 00 kDa triggers cell responses consisting of proautophagy cytokines (IL-10 and TNFα), autophagy, and cell death by apoptosis. These data support a possible mechanism by which extracellular proteins secreted by *C. septicum* can modulate the innate immune response at the infection site, which can serve as a critical pathogenic factor.

One of the characteristic features of myonecrotic diseases caused by clostridia is the absence of phagocytic cells at the site of infection (Junior et al. 2020; Keyel, Heid, and Salter 2011; Ren et al. 2017). This observation suggests an important modulation strategy used by *C. septicum* to evade the innate immune response by preventing the accumulation of phagocytic cells, such as polymorphonuclear cells, monocytes, and macrophages, at the site of infection (Bryant and Stevens n.d.; O’Brien and Melville 2004). It has been shown that certain bacterial cytotoxins, acting alone and more likely together, are capable of producing pores in the membrane of cells of the immune system [2]. Within gram-positive microorganisms, Clostridia exotoxins can induce apoptosis in eukaryotic cells (Ren et al. 2017). However, little information is available regarding which protein or mixture of proteins secreted by *C. septicum* that modulate the apoptotic pathway, and the molecular mechanisms differ from those of gram-negative bacteria. At this stage we only aimed to identify the PPF obtained from C. septicum cultures that cause cell death and to approach to a possible mechanism of immune infection.

The cytotoxicity of the α-toxin of *C. septicum* has been widely studied in different animal cell lines, and it has been shown that its toxic activity varies depending on the cell type used to assess the toxicity mechanism (Kennedy et al. 2009). Moreover, it has been reported that both protoxin and active α toxin exert their cytolytic effect on mammalian cells by forming membrane pores (Hang’ombe et al. 2004), but the involvement of autophagy has rarely been reported. It has been reported that *C. septicum* can secrete, in addition to α toxin, a series of toxins and enzymes that have not been characterized thus far, and thus, their participation in cytotoxicity is unknown. This information is critical for understanding the modulation of innate immune system death mechanism at the infection site. Interestingly, our results indicate that the exposure of mouse peritoneal macrophages to PPF≥100kDa induces apoptosis.

The exotoxins found in this PPF may include *C. septicum* α-toxin; however several authors have reported that the mechanism of cell death induced by this exotoxin does not involve apoptosis, but necrosis. Importantly, α oxin has been found in PPF100-30 kDa (Chakravorty et al. 2015; Cortiñas, Mattar, and Stefanini de Guzmán 1997; Knapp et al. 2010). Our data clearly showed a characteristic DNA ladder fragmentation pattern in macrophages treated with 2.5% PPF≥ kDa. At the same concentration, PPF100-30 kDa caused a broad and random DNA fragmentation pattern consistent with a necrosis cell death mechanism. These results were supported by the morphological pattern of macrophages treated with the same concentration of PPFs. In addition, we found that the expression of the pro-apoptotic protein Bax increased markedly in macrophages treated with PPF ≥ kDa, whereas Bcl-2 expression remained almost unchanged. These results may indicate that the proteins of the PPF≥100 kDa fraction induce apoptosis and that this effect is due to exotoxins other than α oxin.

Our data clearly show that a partially purified fractions of *C. septicum* culture supernatant induce macrophage cell death. The proteins found in the PPF ≥ 100 kDa include various enzymes and toxins that exert cytotoxic effects on host cells (Thomas et al. 2021). Toxins found in this fraction include- lethal toxins such as toxins A and B secreted by *C. difficile*, as well as alpha, beta, epsilon, iota, entero- and type B necrotic- toxins secreted by *C. perfringens* (S. Zhang et al. 2023). Alphatoxin is a well-known virulence factor implicated in gas gangrene in humans and has been shown to cause cell death pathways in host cells (S. Zhang et al. 2023).

Regarding the mechanism of death triggered by exotoxins of molecular weight ≥100 kDa, for example toxins secreted by *C. perfringens* kills fagocytic cells by apoptosis, necrosis, and/or necroptosis (S. Zhang et al. 2023). Likewise, the cytotoxic effect of the *C. histolyticum* culture supernatant has been documented, highlighting the presence of numerous enzymes that can cause the death of the host cell (Jóźwiak et al. 2005). It has been shown that the cytotoxic effects of toxins secreted by clostridium bacteria having a molecular weight ≥100 kDa are due to the presence of various enzymes and toxins that can trigger apoptosis, necrosis or other forms of cell death (Jóźwiak et al. 2005; Navarro, McClane, and Uzal 2018).

On the other hand, incubation of macrophages with low concentrations of *C. septicum* PPF≥ kDa caused autophagy, but the mechanism remains unclear. Autophagy is a lysosomal degradation pathway essential for the cellular response to stressors aimed at preserving cellular survival, differentiation, development, and homeostasis (Bialik, Dasari, and Kimchi 2018; Doherty and Baehrecke 2018). LC3B typically localizes to the cytosol under normal conditions and translocates to autophagosomal membranes when autophagy is triggered (Cuervo et al. 2024). There are two forms of LC3B, LC3B-I and LC3B-II. LC3B-I is the nonlipidated form with a molecular weight of 16L Da, and LC3B-II is the lipidated form with a molecular weight of 14L Da. The conversion of LC3B-I to LC3B-II indicates the formation of autophagosomes (Yu, Chen, and Tooze 2018). Thus, changes in the LC3B-II/LC3B-I ratio are indicative of autophagic activity (Tanida, Ueno, and Kominami 2008). Moreover, the exposure of macrophages to 1% PPF≥ kDa induced autophagy over time (until 4 h). Autophagy is an important process that induces apoptotic cell death in response to *C. difficile* toxin B (Chan et al. 2018; Wang and Cull 2022). In addition, autophagy induced by the PPF≥ kDa could be classified as cell death associated with autophagy, giving an advantage to *C. septicum* during infection .

The balance between pro- and anti-inflammatory cytokines and their effects on cell death pathways is crucial for understanding the pathogenesis of infections and developing potential therapeutic strategies.The detection of TNF-α and IL-10 expressed in macrophages upon infection with *C. septicum* can provide insights into the molecular mechanisms underlying the cell death. The macrophage transcriptomic response to *C. septicum* PPF≥100kDa involves the expression of several survival mRNAs encoding several proteins involved in immunomodulation mechanisms, including autophagy and apoptosis. Among these transcripts, IL-10 and TNFα mRNA (Turner et al. 2014) are critical for triggering autophagy, which can lead to apoptosis. It has been reported that these cytokines induce autophagy and apoptosis in macrophages (Djavaheri-Mergny et al. 2006; Xu et al. 2006); however, the mechanism remains to be fully understood.

However, our data suggest that exposure of mouse peritoneal macrophages to *C. septicum* PPF≥100 kDa can trigger a cell response, including induction of TNFα and IL-10 expression (D.-L. Zhang et al. 2017). Analysis of TNF-α and IL-10 was performed as part of a larger analysis of cytokines secreted by macrophages in response to the toxins contained in the PPF≥100 kDa. The cytokines TNF-α and IL-10 are key mediators in the immune response, where TNF-α generally acts as a pro-inflammatory cytokine and IL-10 functions as an anti-inflammatory/inflammatory cytokine. In the context of infections, particularly those involving *C. septicum*, these cytokines are expressed in macrophages in response to exotoxins produced by the bacteria (Knapp et al. 2010). These cytokines have complex functions in regulating cell survival, inflammation, and immune responses, and can influence processes such as apoptosis and autophagy, which are forms of programmed cell death and cell degradation, respectively (Ge, Huang, and Yao 2018).

TNF-α is known to induce autophagy and apoptosis in several cell types, including mesenchymal stem cells, as part of the inflammatory response (Li et al. 2023). On the other hand, IL-10 is generally considered to inhibit autophagy (Lee et al. 2021). It can counteract the effects of proinflammatory cytokines and may participate in reducing tissue damage during inflammation. Nevertheless, the goal of the study was not to perform a comprehensive analysis of all the cytokine network in the response of macrophages to *C. septicum* exotoxins.

## Conclusions

Our data suggest that exotoxins secreted by *C. septicum* and contained in a PPF ≥ 100 kDa obtained under conditions similar to those found in tissue microenvironments cause inflammation, autophagy and apoptosis in macrophages. Our findings approach the knowledge towards elucidation of a novel mechanism by which exotoxins produced by *C. septicum* deplete innate immune cells, which are critically involved in host defense at the infection site. We are actively working in identifying the protein or mixture of proteins contained in the PPF≥100 kDa that can trigger the inflammatory response and the associated macrophage-cell death mechanism

## Acknowledgments

The authors wish to thank the research support provided by the Universidad Nacional de San Luis (San Luis, Argentina) and the Instituto de Histología y Embriología - Universidad Nacional de Cuyo (Mendoza, Argentina).

## Authoŕs Contribution

Conceptualization, Ortiz Flores RM, Mattar Domínguez MA, and Cortiñas TI; methodology, Ortiz Flores RM, Cáceres CS and Mattar Domínguez MA; software, Ortiz Flores RM; validation, Gomez Mejiba SE and Mattar Domínguez MA; formal analysis, Ortiz Flores RM and Sasso, CV; investigation, Ortiz Flores RM, Cáceres CS, and Sasso CV; resources, Mattar Domínguez MA; data curation, Gomez Mejiba SE, Ramirez DC, and Mattar Domínguez MA; writing—original draft preparation, Ortiz Flores RM and Sasso CV; writing—review and editing, Gomez Mejiba SE, Ramirez DC, and Mattar Domínguez MA; visualization, Gomez Mejiba SE, Ramirez DC and Mattar Domínguez MA; supervision, Cortiñas TI and Mattar Domínguez MA; project administration, Mattar Domínguez MA; funding acquisition, Cortiñas TI and Mattar Domínguez MA. All authors have read and agreed to the published version of the manuscript.

